# Long distance calls: negligible information loss of seabird social vocalisations over propagation down to the hearing threshold

**DOI:** 10.1101/2024.03.04.583271

**Authors:** Anna N. Osiecka, Przemysław Bryndza, Elodie F. Briefer, Katarzyna Wojczulanis-Jakubas

## Abstract

How well does the information contained in vocal signals travel through the environment? To assess the efficiency of information transfer in little auk (*Alle alle,* an Arctic seabird) calls over distance, we selected two of the social call types with the highest potential for individuality coding. Using available recordings of known individuals, we calculated the apparent source levels, with apparent maximum peak sound pressure level (ASPL) of 63 dB re 20 µPa at 1 m for both call types. Further, we created a sound propagation model using meteorological data collected in the vicinity of the little auk colony in Hornsund, Spitsbergen. Using this model, we simulated call propagation up to the putative hearing threshold of the species, calculated to equal ASPL of signals propagated to roughly one kilometre. Those propagated calls were then used in a permuted discriminant function analysis, support vector machine models, and linear models of Beecher’s information statistic, to investigate whether transmission loss will affect the retention of individual information of the signal. Calls could be correctly classified to individuals above chance level independently of the distance, down to and over the putative physiological hearing threshold. Interestingly, the information capacity of the signal did not decrease with propagation. While this study touches on signal properties purely and cannot provide evidence of the actual use by the animals, it shows that little auk signals can travel long distances with negligible information loss. For the animals, this could mean that they can recognize calls of the members of their social networks as far as those calls are actually audible, and support the hypothesis that vocalisations could facilitate long-distance communication in the species.

## Introduction

The ability to recognise one’s social partner – e.g. offspring, mate, or neighbour – is necessary to maintain stable social bonds. Colonial animals, such as seabirds, often rely on vocal cues to find each other in crowded aggregations (e.g., Klenova et al. 2012, Favaro et al. 2015, 2016, Calcari et al. 2021, Bowmaker-Falkoner et al. 2022). But how reliable is such communication at a distance?

While under some conditions, acoustic signals can travel over extreme distances (e.g. a blue whale’s song theoretically travelling through the oceans), this is not always the case. The propagation of a soundwave, i.e. how it moves through and changes in an environment, depends on a number of factors. First of all, signals of lower amplitudes will degrade much faster due to spherical spreading, than louder sounds. Additionally, as the sound propagates, its higher frequency content will be gradually filtered out, leaving only the lower frequency components at larger distances (and finally filtering these as well). How exactly this filtering will occur, and how fast will a soundwave travel, will be impacted by the medium in which it is travelling – its density, humidity, pressure, and more. At some point, a signal’s amplitude will be so low, and/or its frequency content so degraded, that it will no longer carry the information first encoded in it by the sender – and of course, as a result, the receiver will not be able to decode it.

Little auks (*Alle alle*) are highly colonial seabirds navigating complex social networks (Wojczulanis-Jakubas *et al*. 2022). Little auks are also very vocally active (Osiecka *et al*. 2023a), and their calls can carry a richness of static (Kidawa *et al*. 2023, Osiecka *et al*. 2023b, 2024a) and dynamic (Osiecka *et al*. 2023a, 2024b) information. The most complex call of the little auk repertoire, the *classic* call, is a long, compound signal with apparent formants, composed of a series of three types of syllables (Fig. 1; Osiecka *et al*. 2023a). It is a social call produced in a range of contexts, both by animals sitting inside their rocky nest chambers, and in flight, e.g. by birds returning to the colony from the foraging grounds (Osiecka *et al*. 2023a). While it carries no information on the caller’s sex or size, nesting partners tend to match certain properties of their *classic* calls (Osiecka *et al*. 2023b). This vocalisation carries reliable information on the sender’s identity, mostly within its spectral centre of gravity, fundamental frequency, duration, amplitude modulation rate, and frequency variation (in this order; Osiecka *et al*. 2024a), and has a higher information capacity than any other call type of the species (Osiecka *et al*. 2024a). The *classic* call likely plays a role in long-distance communication, possibly facilitating coordination of social behaviour. Therefore, it is likely to remain stable over behaviourally useful distances.

**Figure 1.**
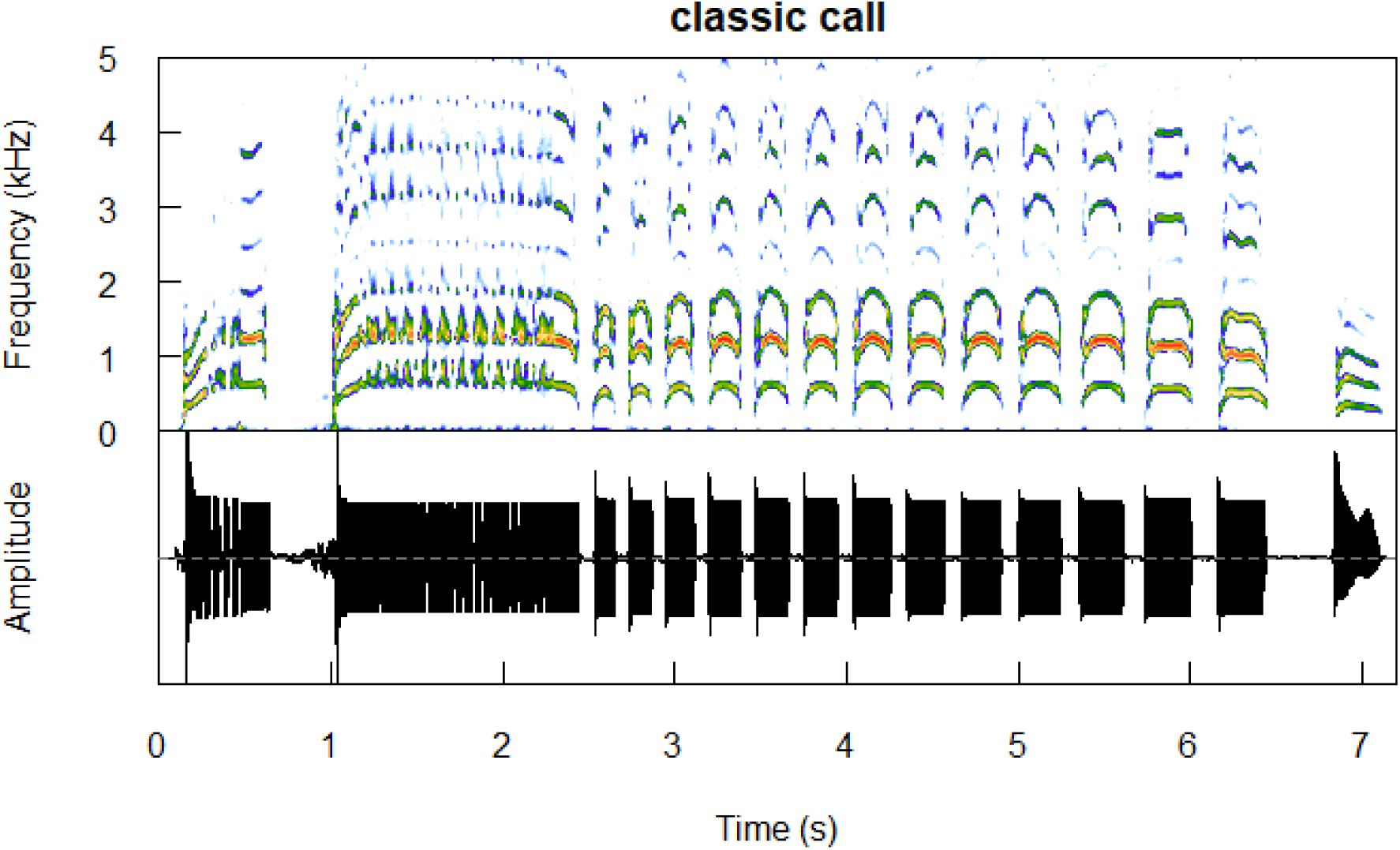
A sample *classic* call produced by an adult male (ring no. DA48567). Spectrogram plotted using the *seewave* package (Sueur *et al*. 2008).

Another social call emitted in a range of situations, and both inside the nest and in flight, is the *single* call; a brief, one-syllable vocalisation (Fig. 2; Osiecka *et al*. 2023a). Like all little auk call types, this call is highly individually specific, and can be classified to an individual with the highest precision among all call types (Osiecka *et al*. 2024a). While the exact function of this call remains unknown, due to its short duration (less than 0.5 s; Osiecka *et al*. 2023a) and simple structure, it can be expected to serve in short-distance communication.

**Figure 2.**
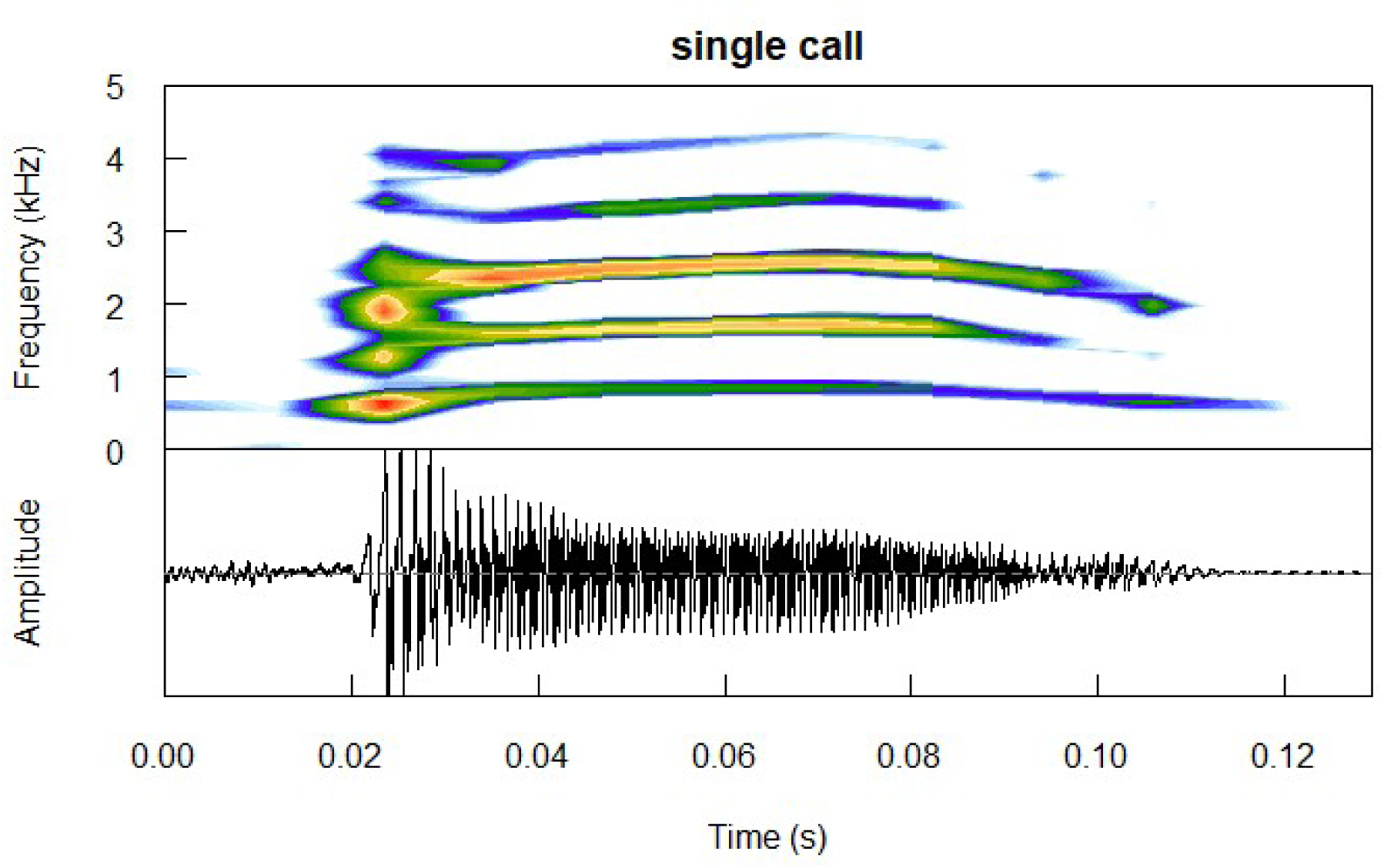
A sample *single* call produced by an adult male (ring no. DA48567). Spectrogram plotted using the *seewave* package (Sueur *et al*. 2008).

Here, we investigated how well the identity information encoded in the *classic* and *single* calls, which are both used by little auks throughout the entire breeding season, is maintained over distance, purely from a signal perspective, i.e. the transmission-related changes to carrying capacity of the vocal communication channel. We expected the *classic* calls to propagate better, over longer distances than the *single* calls, which likely serve short-distance communication. To test this, we created a theoretical propagation model using local meteorological data, and, using sample calls of the two aforementioned types recorded from known individuals, we simulated call propagation down to the putative hearing threshold. We then investigated the information content of those propagated calls.

## Methods

All analyses were performed in Python v. 5.11 (Rossum and Drake 1995) and R environment (v. 4.1.3), and full codes together with raw data have been provided in the supplementary materials (see Data availability statement). Visualisations use scientific colour palettes (Cramieri 2018 ; Cramieri *et al*. 2020) from package *khroma*.

### Study site and subjects

This study used previously published acoustic recordings (see detailed description below; Osiecka *et al*. 2024a). These recordings were collected during fieldwork in Hornsund, Spitsbergen, Norwegian High Arctic, over the incubation period in 2019-2020, under permit from the Governor of Svalbard (20/00373-2). This included handling (e.g. colour-ringing and measuring) the birds for standard ornithological procedures by a licensed ringer (KWJ, permit no. 1095, type: C, issued by Museum Stavanger, Norway), in order to be able to identify the focal individuals (see description below in the *Acoustic data* section). This study focused on 18 nesting pairs, i.e. 36 birds in total.

The study colony in Hornsund is comprised of the lower: 59-90 m a.s.l. and upper plot of the colony: 122-172 m a.s.l. Little auks maintain their flight height above their colony plots, and only descend for landing. For the purpose of this study, we selected 100 m as a representative flight height for the lower plot, i.e. the animals recorded in this study. This choice to select a flight height lower than the upper plot was made as a conservative measure to avoid accidentally increasing the modelled active space of little auk sounds (see model details below).

### Acoustic data

Audio material was collected via an Olympus ME-51S stereo microphone (-40 dB sensitivity at 1 kHz, frequency response 100-15,000 Hz +/-3 dB) placed inside each nest (a rock crevice/chamber, with floor covered with pebbles, Wojczulanis-Jakubas *et al*. 2022) at approximately 10 cm from the birds inside, in such a way as to not disturb the birds’ normal activities. Each microphone was connected to an Olympus LS-P4 digital voice recorder (sampling rate 48 kHz, 16 bits, high gain) placed outside of the nest chamber and hidden under a rock to prevent both disturbance of the animals and damage to the equipment. Each nest was recorded three times over the incubation period, with recording sessions lasting 48 h and spaced about equally in time (i.e., around eight days in between recording sessions).

Sound recordings were paired with video monitoring of the nest entrance, so that we could see the birds entering and exiting their nesting chambers and extract the times at which only one known (ringed with a unique colour code) individual was present inside the nest chamber. Audio recordings from those periods were manually processed, resulting in the acoustic database of vocalisations produced by known individuals inside the nest. For more details on the field procedures, refer to Osiecka *et al*. 2024a.

### Apparent sound pressure level

To calculate the real-life sound pressure levels from the collected recordings, we first calibrated the equipment. First, a class II sound level meter (Volcraft SL-451) was calibrated with a class II sound level calibrator (Volcraft SLC100) following instructions provided by the producer. Then, a 1 kHz tone was played using a JBL Flip 5 loudspeaker placed at 1 m from the recorder and sound level meter, and recorded with the same equipment and set-up as used in the field recordings. The obtained recording was used in end-to-end calibration of all digital audio recordings in Raven Pro 1.6.5, following the software specifications (https://ravensoundsoftware.com/knowledge-base/calibrating-recordings-in-raven-pro/).

Back-calculated sound pressure levels are termed *apparent sound pressure levels* (hereafter ‘ASPL’, Møhl *et al*. 2000). ASPL (dB rms re. 20 µPa) at 10 cm of each vocalisation was extracted in Python using *numpy* package to obtain peak (i.e. the highest absolute magnitude of the signal) and root-mean-square (RMS, i.e. the RMS amplitude over signal duration, using the 95% energy threshold criterion; Madsen and Wahlberg 2007) values. The ASPL at 1 m, i.e. the Source Level (SL), was calculated as:

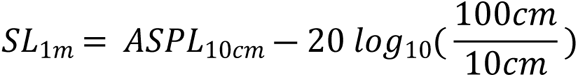

To estimate a global mean of the ASPL values at 1 m, we first calculated the mean ASPL value for each individual, followed by a population mean. This was done for both call types, with peak and RMS values used separately. The obtained mean values were then compared between the call types using Welch two sample t-test (function *t.test*).

### Meteorological data

Long-term geosystem monitoring data are publicly available from the Polish Polar Station in Hornsund, Institute of Geophysics, Polish Academy of Sciences (https://monitoring-hornsund.igf.edu.pl). For the purpose of this study, we selected data from 1983-2021, for which full meteorological information was available (as per August 2023, when the analysis was performed), focusing on May-August, i.e. the breeding period of the little auk (Wojczulanis-Jakubas *et al*. 2022). Because those months are characterised by very different mean temperature, pressure, and relative humidity values (Fig. 3) – and therefore different sound attenuation properties – we considered each month separately, using the 40-years average of each month in the following analyses.

**Figure 3.**
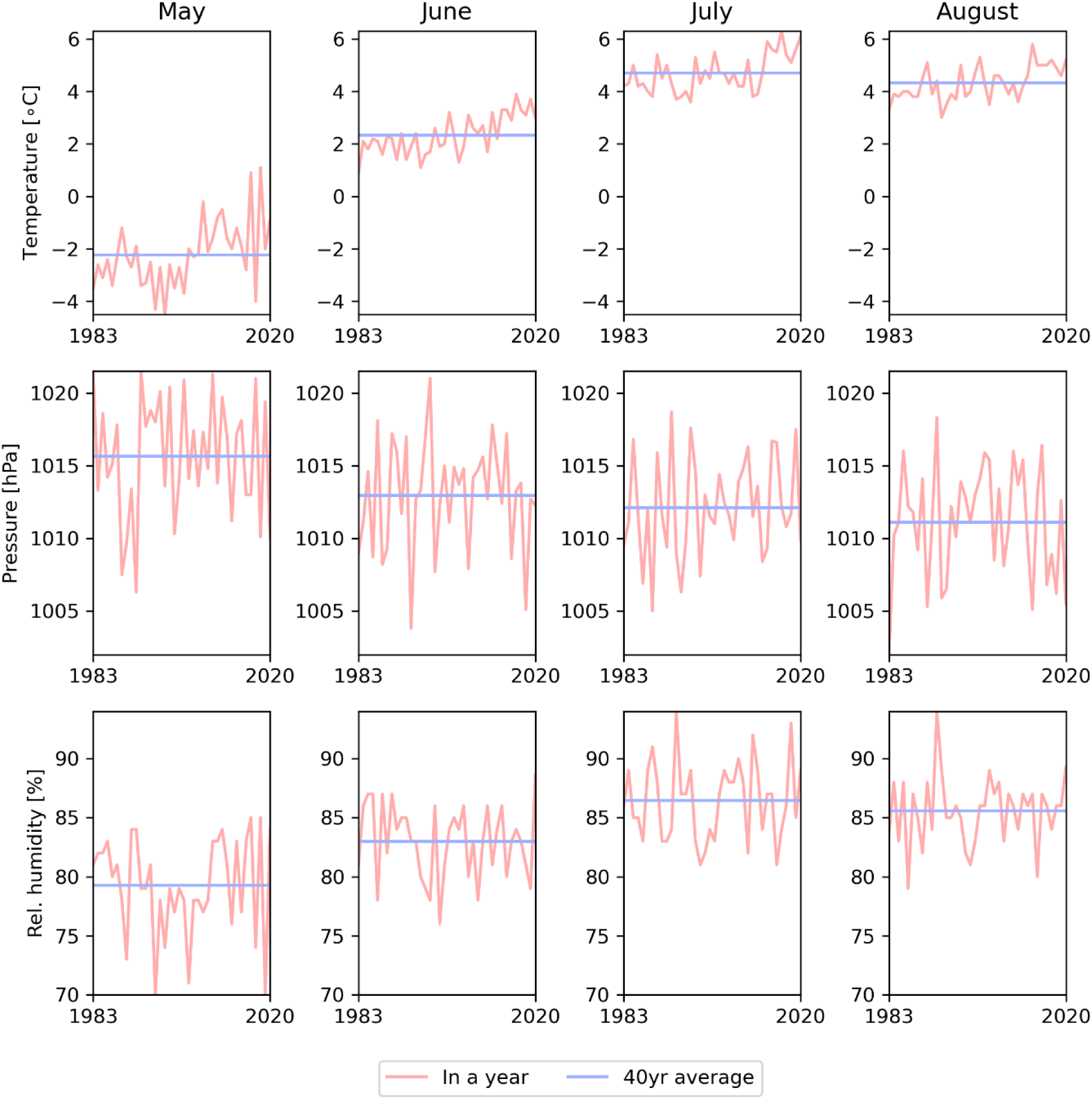
Annual (pink) and 40-year average (blue) meteorological data from Hornsund over the little auk’s breeding period (May-August).

### Propagation model

To model propagation of signals over distance, we used a spherical spreading model with the atmospheric absorption factor α based on the ISO 9613–1 standard (ISO 1993). The spherical spreading model describes how the energy of different frequency components of the signal changes over distance, working somewhat as a low-pass filter (i.e. the energy content of higher frequencies is lost earlier over propagation).

Note that this model comes with necessary simplifications: that is, it assumes simple spreading in perfect conditions, i.e. without added noise, in the absence of wind, and excluding excess attenuation. Simple spherical spreading was chosen based on the following: (1) We decided to model propagation of calls produced in flight, and not in the nest, to simplify the model. Therefore, the signal source is an individual bird in flight, that is roughly 100 m over ground. This model is hence simplified to omit the impact of local topography on sound propagation (see Guibard *et al*. 2022). While the *classic* and *single* calls are frequently produced both in flight and inside the nests, note that the calls used here were recorded inside the nest, since this is the only way we could control for the birds’ identity. The implications of this are addressed in the Discussion; (2) The Hornsund ornithogenic tundra is an open habitat with a dense vegetation cover composed of species reaching a maximum of approximately 20 cm in height (Zmudczyńska *et al*. 2009), and is therefore expected to minimally degrade acoustic signals (Hardt and Benedict 2020); (3) The dense vegetation cover creates a soft substrate, so contribution of reflections is expected to be minimal; and (4) diel variations in meteorological conditions during the Arctic day are dictated by sea ice conditions rather than time day-night cycles (Osuch and Wawrzyniak 2017), which means that reflections from different layers of the atmosphere are also expected to be minimal.

The ISO 9613-1 standard gives fitted equations for atmospheric attenuation α as a function of frequency that is dependent on temperature, pressure and relative humidity of the air. The model is valid at altitudes below 10’000 meters, and so well within our case. As described above in the Meteorological data section, we used the local mean monthly values of relevant parameters, and subsequently α was calculated on those mean monthly values. We used the average values of the entire monitored period (1983-2021) rather than climate change-related patterns, since there was no apparent change in sound attenuation properties over the decades (Fig. 4).

**Figure 4.**
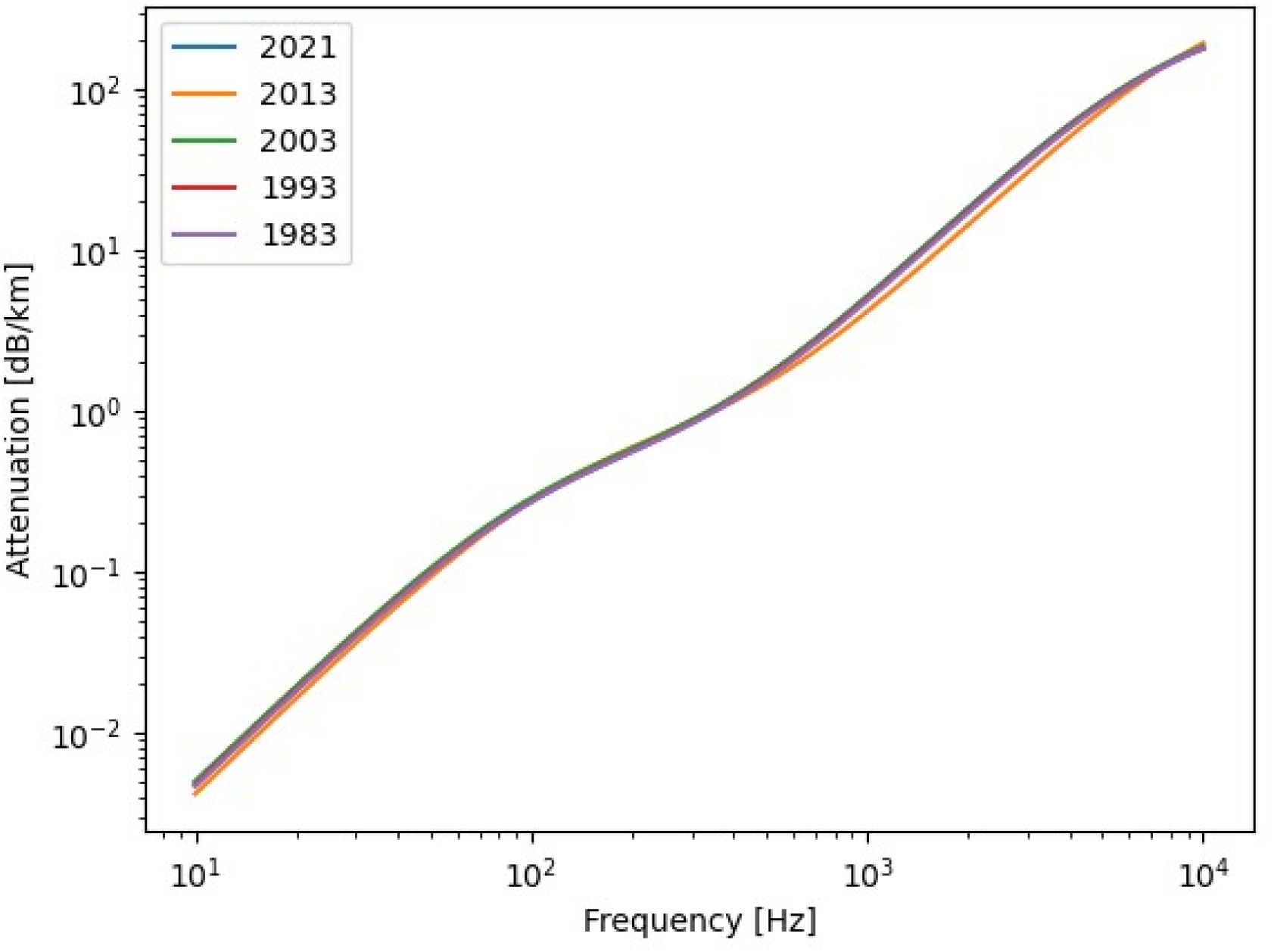
Sound attenuation at different frequencies, calculated from mean May conditions in Hornsund over the monitored period 1983-2021, based on the ISO 9613-1 standard. There is no apparent shift in attenuation profiles over the years.

The resulting spherical spreading model is given by the following equation:

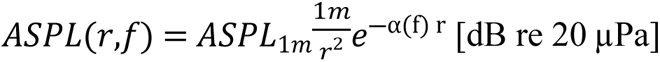

Where *e^x^* is the natural exponential function, r is the distance (in metres), and α is a function of frequency as per ISO 9613-1. The full code of the propagation model is available in the Supplementary Materials.

### Choice of the propagation distances

Since there is currently no information available on the hearing thresholds of the little auk, we used the in-air auditory measurements of another, related diving alcid species, the Atlantic puffin (*Fratercula arctica*), as a reference. The average physiological hearing threshold (measured using auditory evoked potential methods) in the alcids seems relatively similar across species, namely down to 10-20 dB re 20 µPa in the 1-2.5 kHz frequency range for the Atlantic puffin (Mooney *et al*. 2020), down to 13 dB re 20 µPa in the 1-3.5 kHz range in the common murre (*Uria aalge*; Smith *et al*. 2023a), and down to 17 dB re 20 µPa in the 1-3.5 kHz range for the marbled murrelet (*Brachyramphus marmoratus*; Smith *et al*. 2023b). We chose 1000 m as the maximum propagation distance, with calculated ASPL at this distance roughly corresponding to the minimum physiological hearing threshold (i.e. the lowest SPL within the studied frequency range that still elicited brain activity during experimental procedures) of the Atlantic puffin (Mooney *et al*. 2020; Fig. 5).

**Figure 5.**
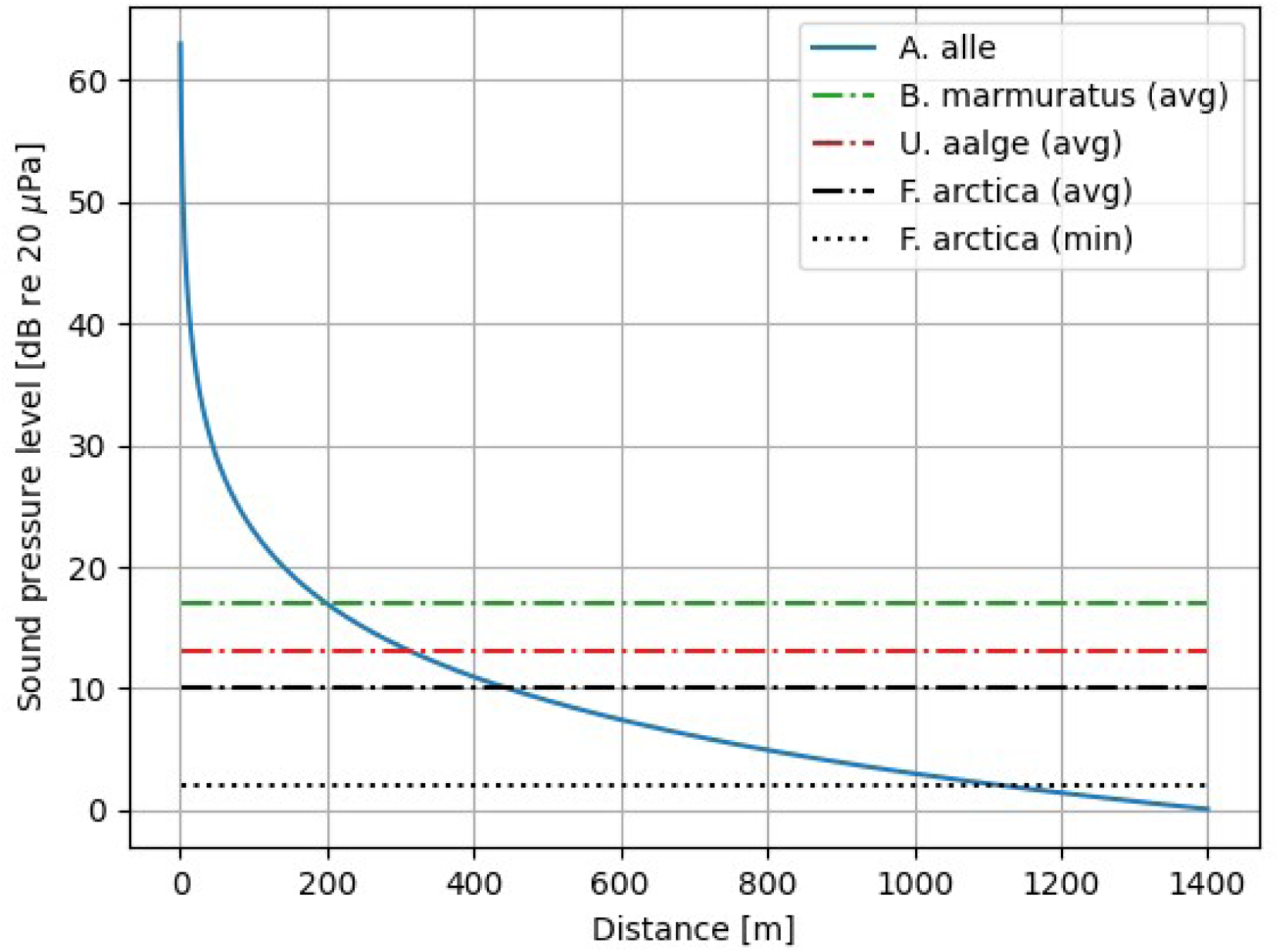
Average (broken lines) and minimum (dotted line) physiological hearing thresholds of three alcid species allow us to predict the expected distances at which the little auk vocalisation can still be heard (based on simple spherical spreading, i.e. -6 dB re 20 µPa per doubling of distance).

The propagation model used calibrated recordings of known individuals at 10 cm as input files. Each file was propagated (i.e., modelled in meteorological conditions for May-August separately) to 1, 2, 4, 10, 21, 46, 100, 215, 464, and 1000 m (from here on, 1-1000 m), creating a separate audio file as an output. In other words, each original call was propagated to 10 distances in mean conditions of four separate months, that is 40 times in total. Note that this does not mean performing actual propagation experiment in the air, but purely mathematical modelling resulting in selectively filtered vocalisations.

### Acoustic analysis

All obtained (i.e. propagated) audio files were batch-processed in R, using the *soundgen* package (function *analyze* with settings adjusted to the little auk: dynamic range = 60 dB, pitch floor = 500 Hz, pitch ceiling = 2000 Hz, step = 5 ms) to extract a set of 15 acoustic parameters (Table 1). Both raw audio and the resulting analysed datasets can be found in the supplementary materials.

**Table 1.**
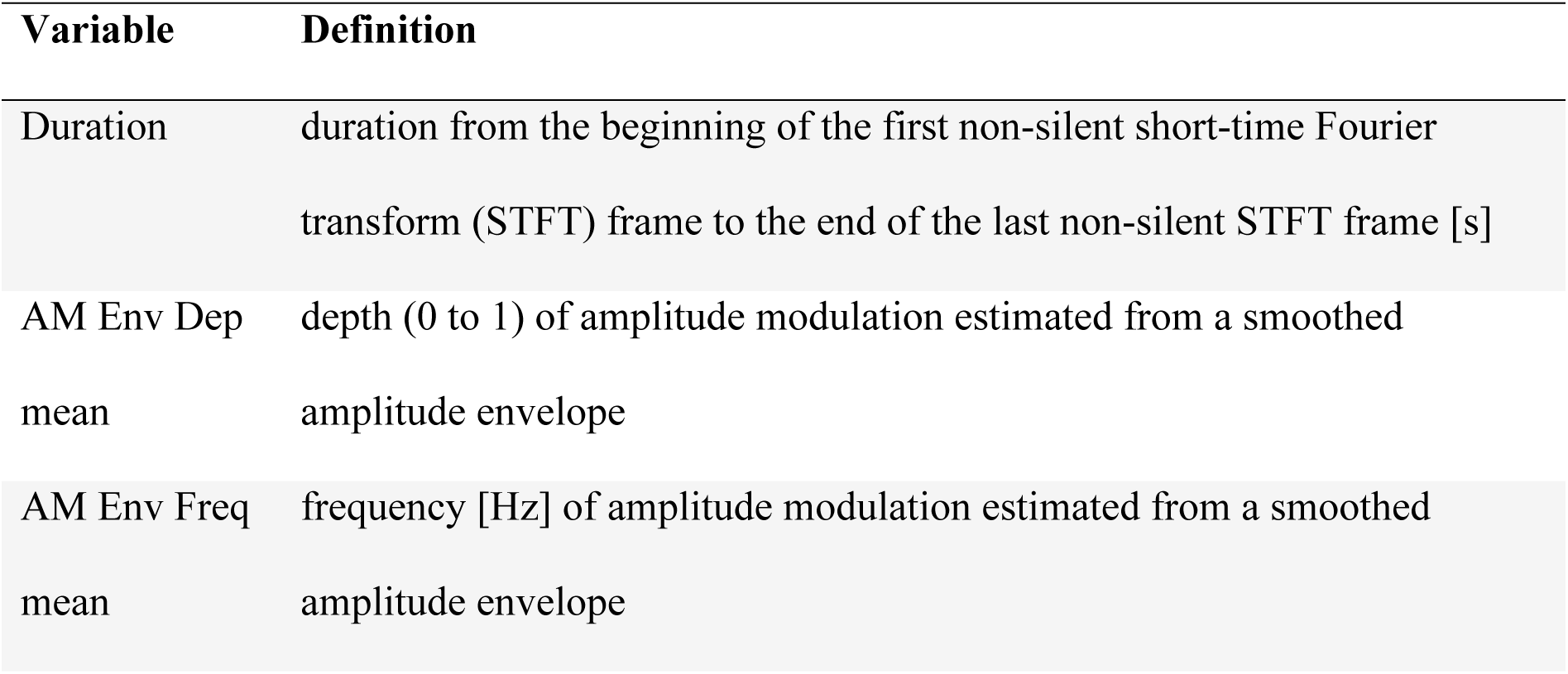
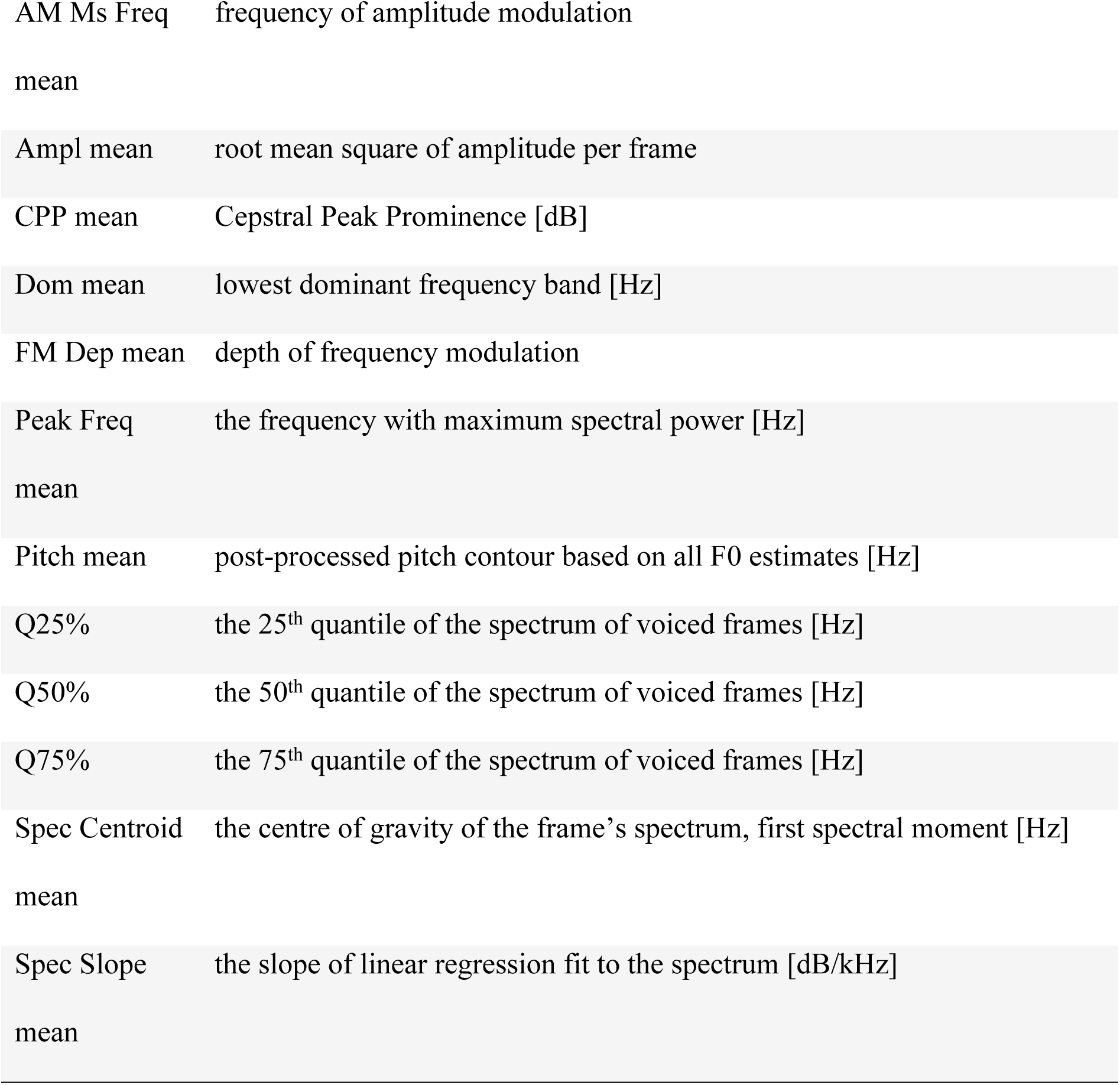
Raw acoustic parameters extracted from audio files. Variable explanations as per *soundgen* package.

The dataset was first cleaned, i.e., entries with missing values (that is, raw acoustic parameters that could not be correctly extracted) removed. We also reduced the dataset to the individuals with at least 200 entries (i.e., at least five calls propagated four times to 10 distances). This reduced the dataset to 5521 *classic* call entries from 11 individuals, and 2640 *single* call entries from six individuals.

To reduce data dimensions for further analyses, this cleaned dataset was subsequently tested for Kaiser-Meyer-Oklin factor adequacy (function *KMO,* package *EFAtools;* Supplementary Table 1), and then used in a Principal Components Analysis (PCA; function *prcomp*, package *stats;* Supplementary Tables 2 and 3). This was done separately for each of the two call types.

### Classification to individual over distance

To check how well can propagated calls be classified to the caller independently of the distance, we performed the following analysis. For each call type, we selected the principal components with eigenvalues > 1 (Supplementary Table 2) as input variables. The corresponding PC scores of all obtained calls (i.e. calls propagated at distances 1-1000 m) for which we were able to extract the full set of acoustic parameters specified in Table 1 were used in a permuted discriminant function analysis (pDFA; Mundry and Sommer 2007), to see how well can calls be classified to the caller independently of the distance. This pDFA was conducted in a nested design, using the *pDFA.nested* function (R. Mundry, based on function *lda* of the *MASS* package), on all available calls (5521 for the *classic* call, and 2640 for the *single* call) of all the subjects (11 for the *classic* call, and six for the *single* call). Since the same calls were propagated in conditions corresponding to the four focal months (May-August), we used the file name as a control factor to correct for multiple sampling. We ran a total of 1000 permutations for the analysis. This was done separately for the two call types, for all distances pooled together and each distance separately.

Furthermore, to see how well calls propagated to different distances cluster to individuals, we performed a set of additional analyses using support vector machine (SVM) classifiers. First, to establish the approximate number of nearest neighbours to use, we used the *kNNdistplot* function of the *dbscan* package (Hahsler *et al*. 2019). We then reduced the data dimensions of the raw, cleaned datasets using supervised uniform manifold approximation and projection (S-UMAP; *uwot* package, *umap* function), with minimum distance = 0.5, n_neighbours = 500 (*classic*) or 200 (*single*), using the Euclidean metric. This gave us two-dimensional coordinates, subsequently introduced to the SVM classifiers. The data were first subset into distances, and subsequently into 8:2 training:test datasets. A classification task was built for each subset (*mlr* package, function *makeClassifTask* with individual ring number as target). A learner was then created using *makeLearner* function of the *mlr* package, and corrected for individual weights due to the uneven sampling of different individuals (*mlr* package, *makeWeightedClassesWrapper* function). The weighted learner was then trained (*mlr* package, *train* function) on the training task, and used to classify the task (*mlr* package, *predict* function). Classification accuracy of the SVM was extracted using the *performance* function of the *mlr* package. The accuracy was then compared in a simple linear model (function *lm*). This was performed for each call type and propagation distance separately.

### Information loss over distance

To investigate the possible loss of information content of the signal over distance, we used Beecher’s information statistic, *H_s_*(Beecher, 1989), which informs about the information capacity of a signal. To calculate *H_s_*, we used all PC scores into the *H_s_*calculation (function *calcHS, IDmeasurer* package). This was performed on subsets of calls propagated at different distances (1-1000 m, 10 calculations per call type in total).

## Results

### Apparent sound pressure level

The apparent sound pressure levels, expressed as the mean peak ASPL and mean ASPL RMS, were slightly higher for the *classic* than *single* calls (Table 2). However, the maximum peak ASPL and mean ASPL RMS were similar for the two call types (Table 2).

**Table 2.**
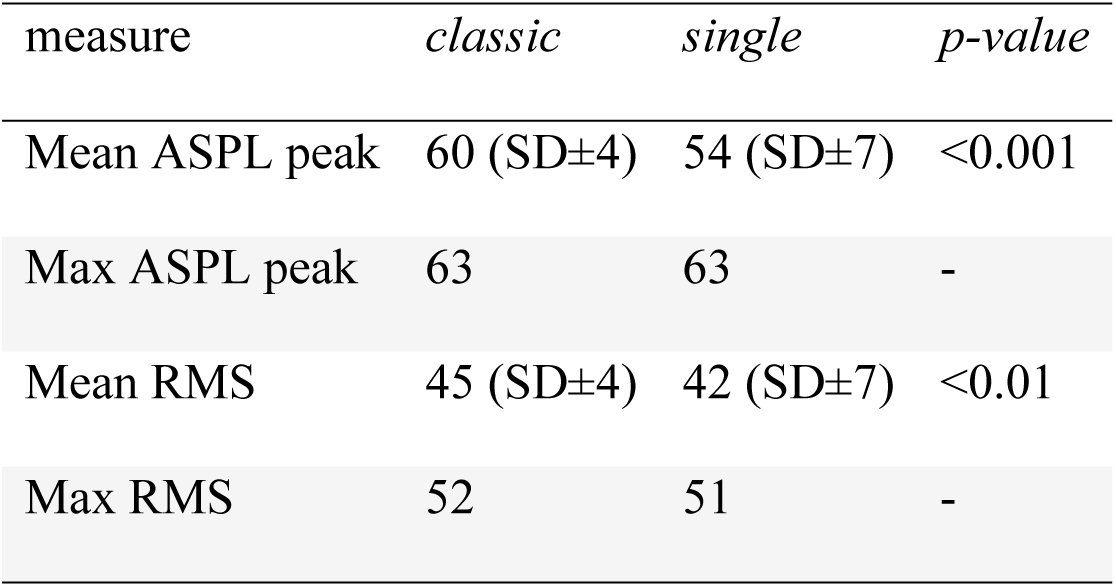
Maximum and mean SL values of the call types. All SL values are given in dB re 20.

### Classification to individual over distance

Call structure remained stable over large distances (Figs. 6-7), and calls could be classified to the correct individual above chance levels independently of the distance (Tables 3-4). Clustering accuracy did not decrease with distance (Figs. 8-9, Table 5).

**Figure 6.**
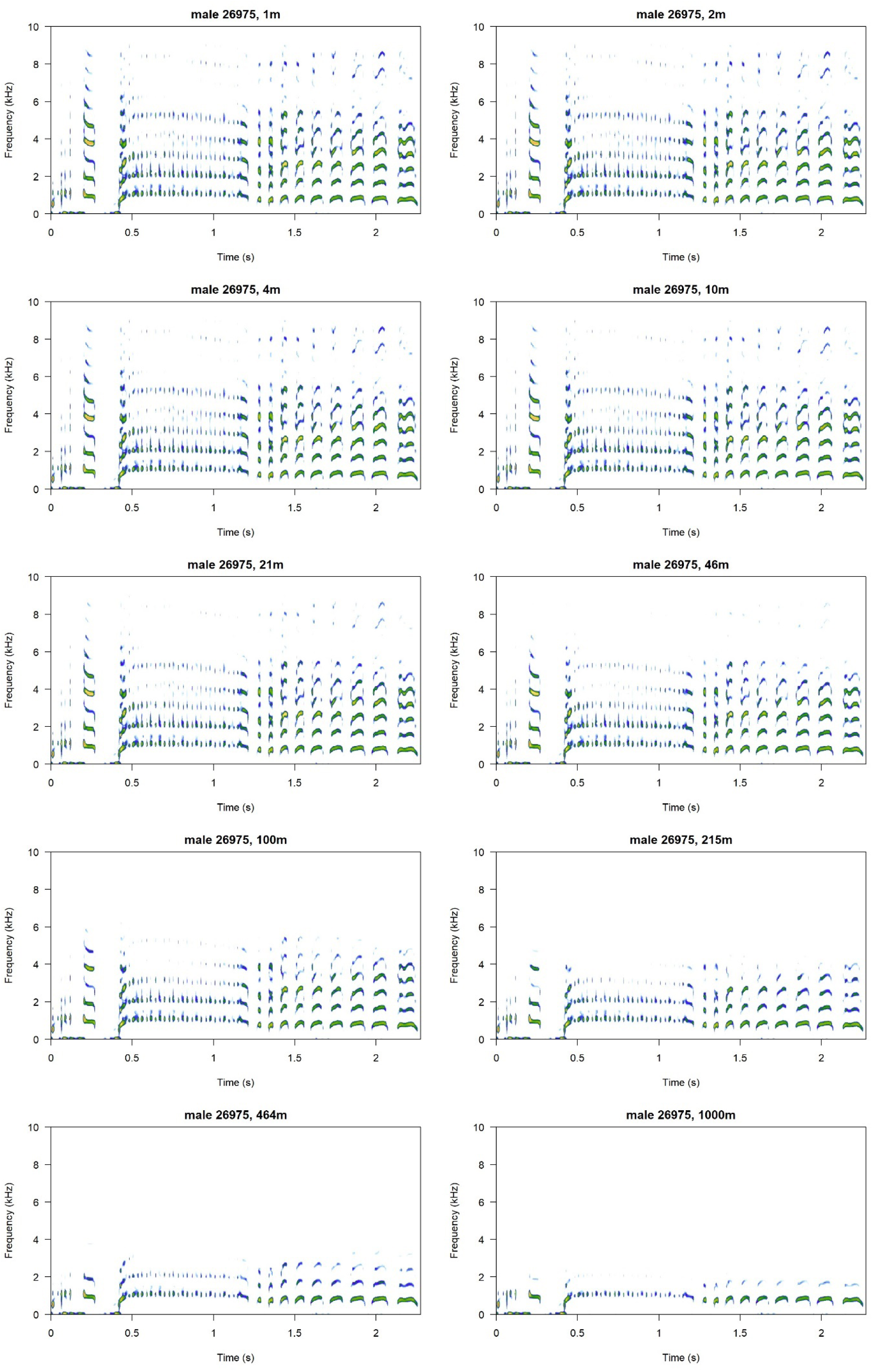
A sample *classic* call propagated at 10 exponential distances in a range of 1-1000 m. Notice that the signal remains very stable across the distances, and harmonics are only lost at extreme distances, close to the putative physiological hearing threshold. Spectrograms plotted using the *seewave* package (Sueur *et al*. 2008).

**Figure 7.**
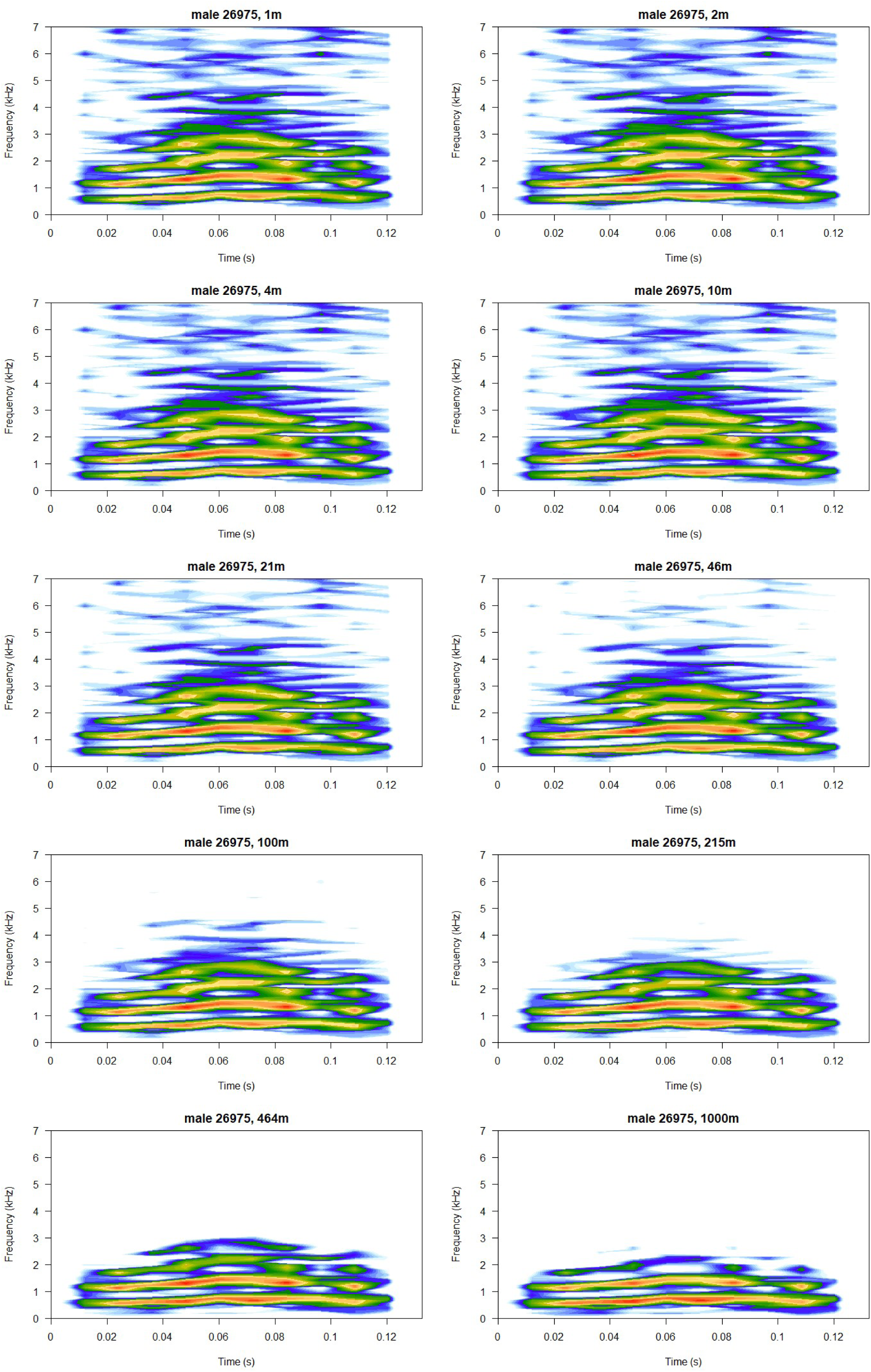
A sample *single* call propagated at 10 exponential distances in a range of 1-1000 m. Notice that the signal remains very stable across the distances, and harmonics are only lost at extreme distances, close to the putative physiological hearing threshold. Spectrograms plotted using the *seewave* package (Sueur *et al*. 2008).

**Fig 8.**
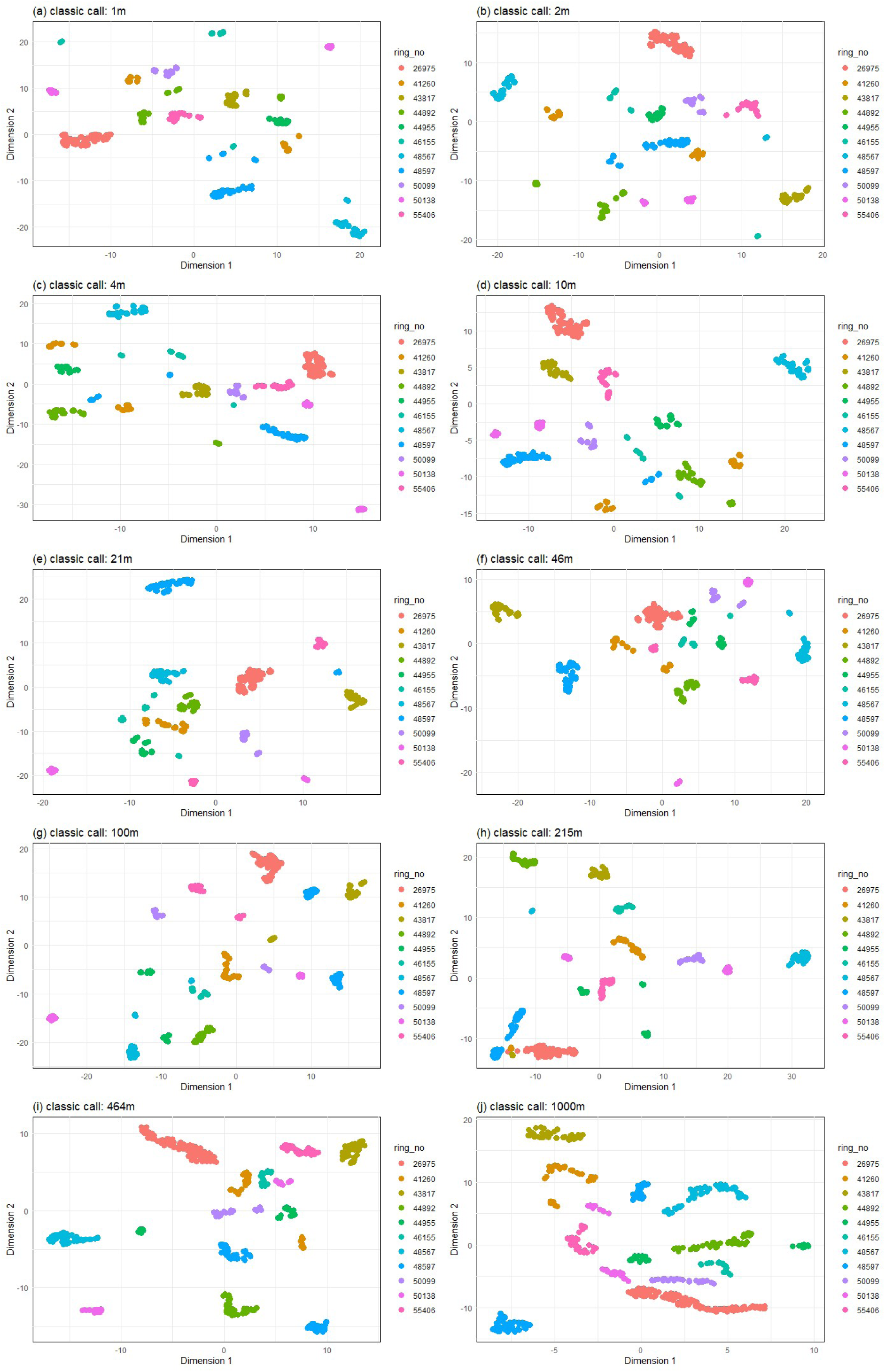
S-UMAP classification of the *classic* call to individual over 10 different propagation distances.

**Figure 9.**
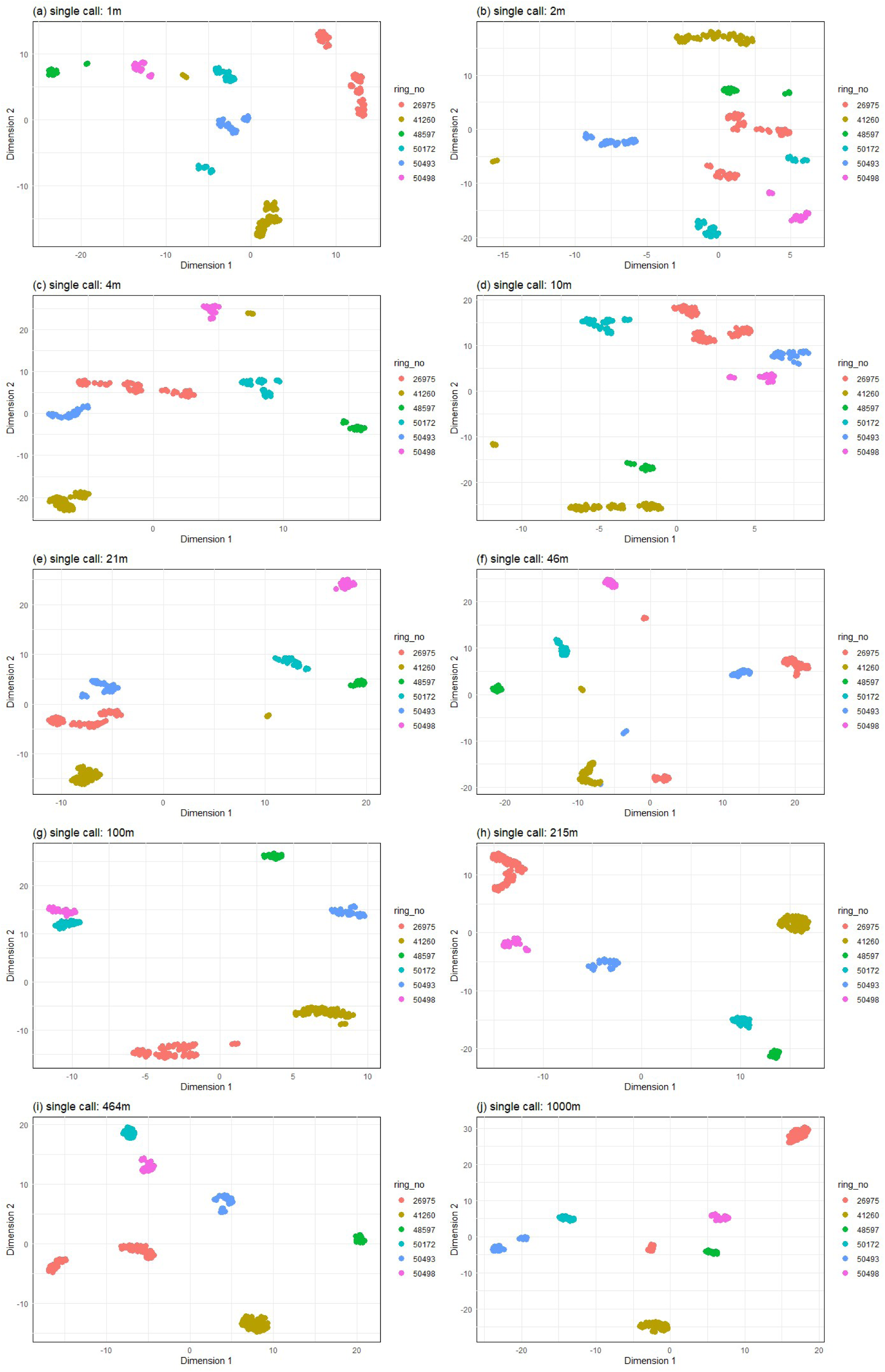
S-UMAP classification of the *single* call to individual over different propagation distances.

**Table 3.**
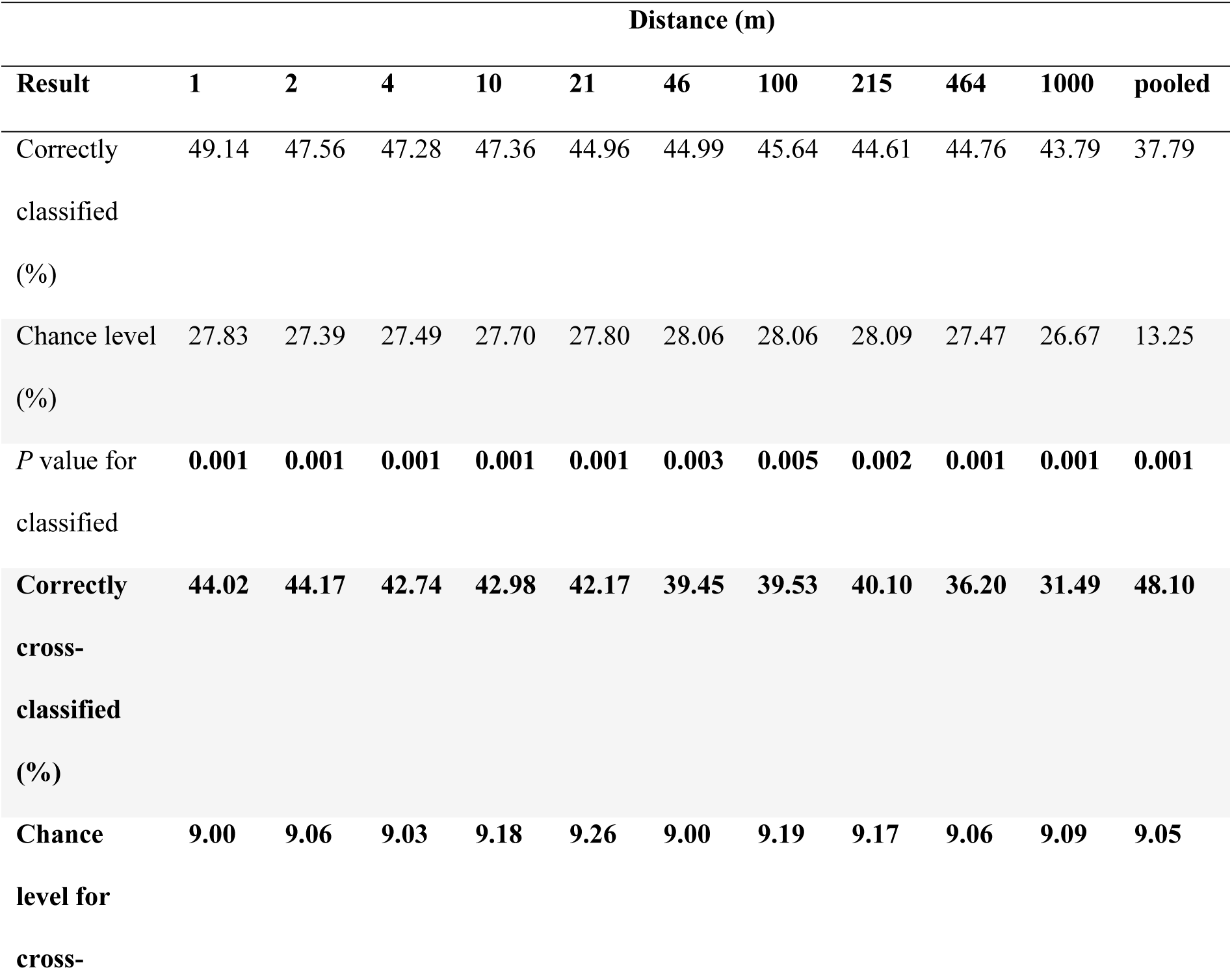
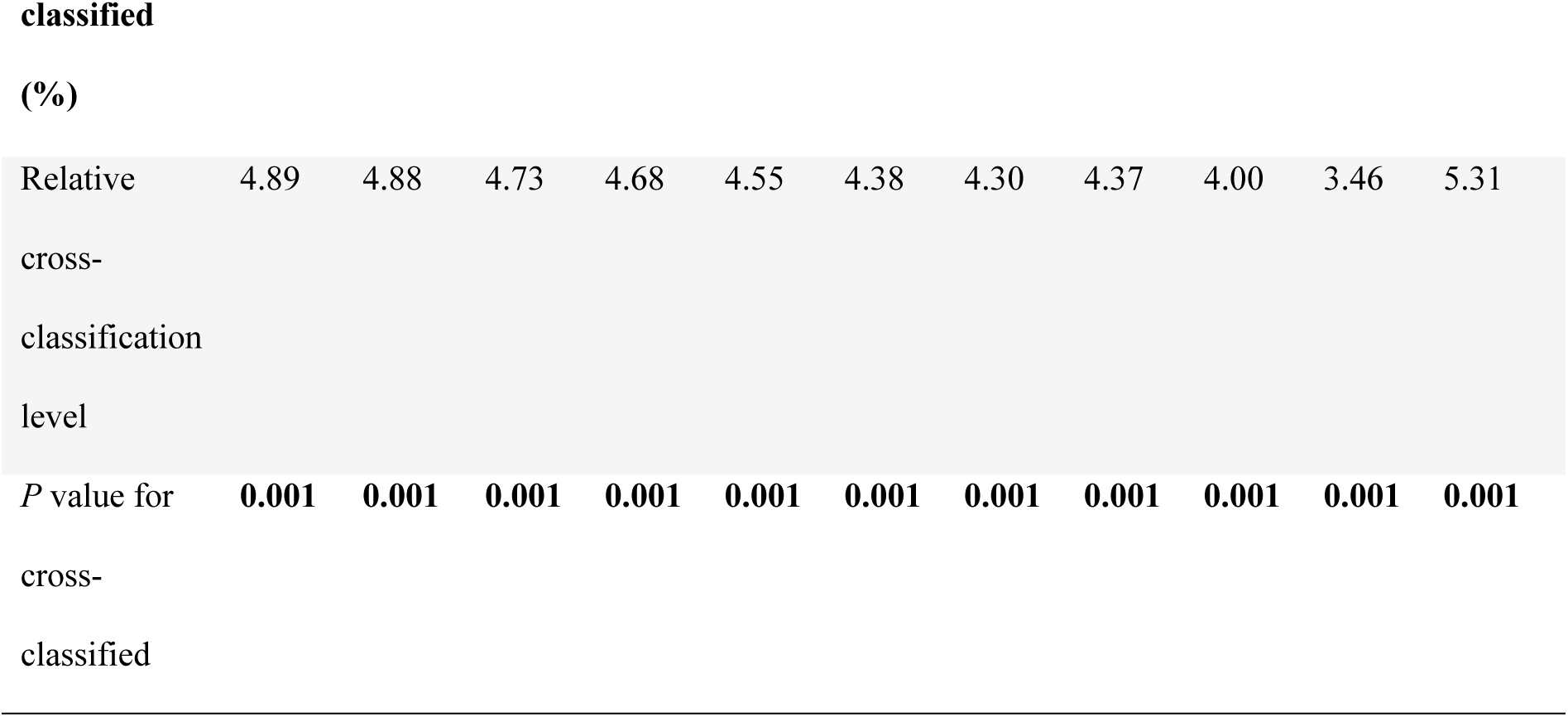
Results of the permuted discriminant function analysis for *classic* calls propagated at distances from 1 to 1000 m (552 calls of 11 individuals per distance), as well as for all distances pooled together (5520 calls of 11 individuals), using the principal components of eigenvalues 1. Calls could be reliably classified to individuals above chance levels independently of the distance.

**Table 4.**
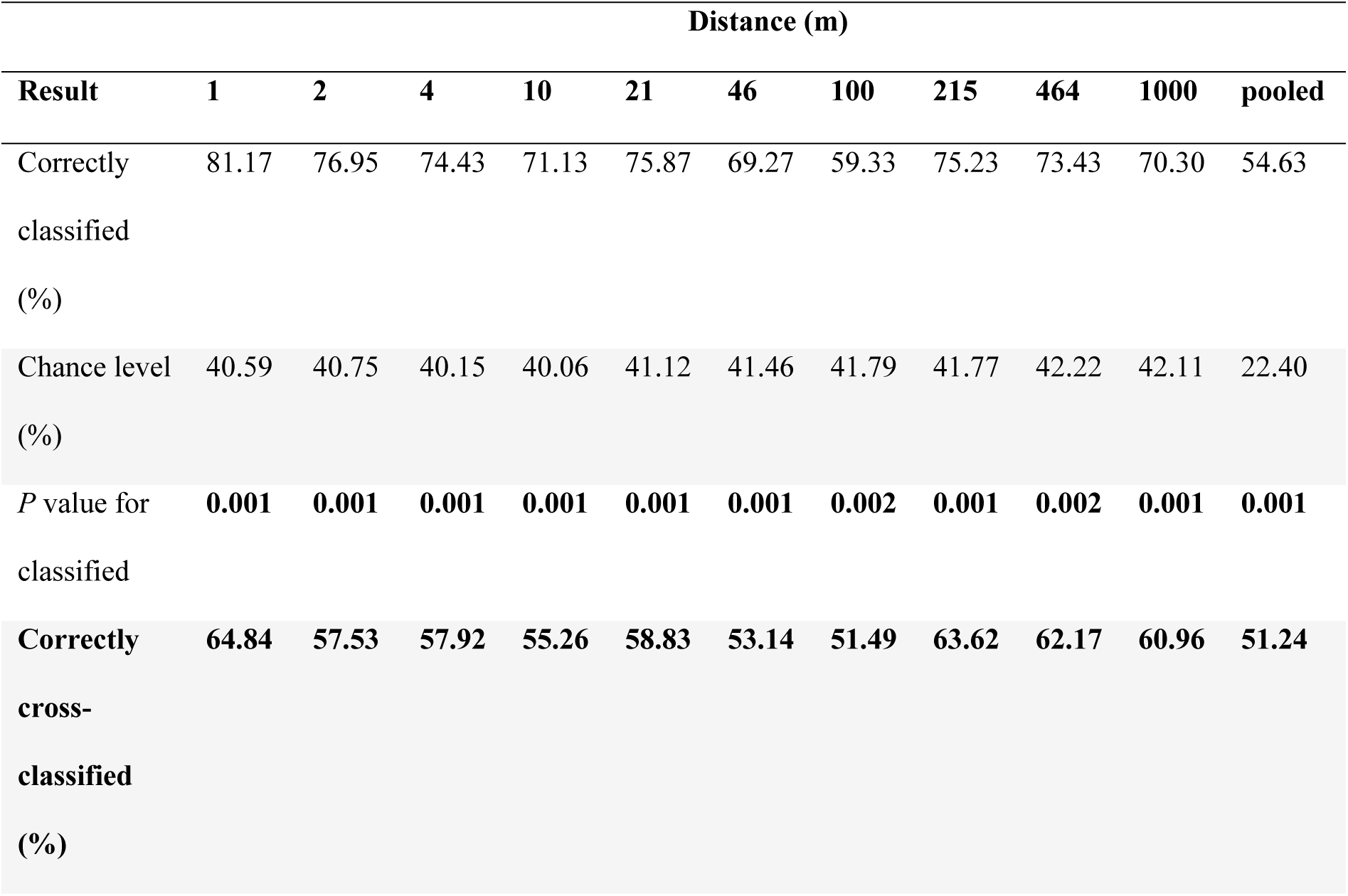
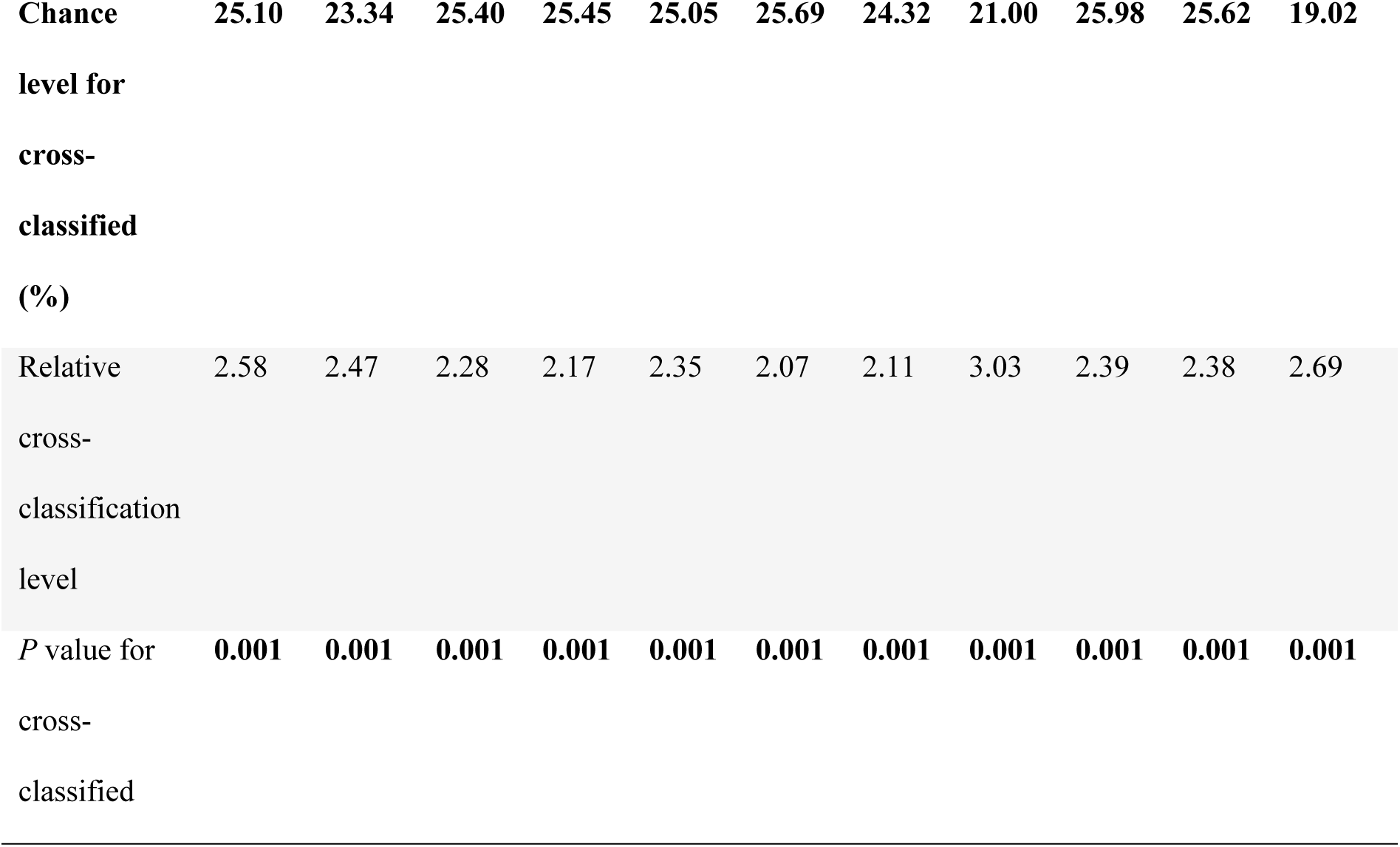
Results of the permuted discriminant function analysis for *single* calls propagated at distances from 1 to 1000 m (264 calls of six individuals per distance), as well as for all distances pooled together (2640 calls of six individuals), using the principal components of eigenvalues 1. Calls could be reliably classified to individuals above chance level independently of the distance.

**Table 5.** Accuracy of classification to individual using SVM based on S-UMAP reduced data. Distance Accuracy [%].

| Distance | Accuracy [%] |  |
| --- | --- | --- |
|  | *classic* call | *single* call |
| 1 | 58 | 73 |
| 2 | 62 | 72 |
| 4 | 61 | 85 |
| 10 | 61 | 69 |
| 21 | 59 | 74 |
| 46 | 66 | 78 |
| 100 | 56 | 83 |
| 215 | 62 | 77 |
| 464 | 57 | 89 |
| 1000 | 65 | 72 |
| p-value | 0.4 | 0.9 |

### Information loss over distance

The information capacity of the *classic* call did not decrease with distance, theoretically allowing for a distinction of essentially constant number of individuals as at the source (Table 6). By contrast, the *short* call seemed to be particularly individually specific at a very short range (1 m), and maintained roughly 50% of its original information content over propagation.

**Table 6.** Beecher’s statistic’s values in the propagated signals. Column *meaning* specifies how many individuals can be theoretically distinguished based on the signal alone.

| *classic call* |  |  |  | *single call* |  |  |
| --- | --- | --- | --- | --- | --- | --- |
| Distance [m] | $H_s$ significant | $H_s$ all | meaning | $H_s$ significant | $H_s$ all | meaning |
| 1 | 2.8 | 2.8 | 7 | 3.5 | 3.6 | 12 |
| 2 | 2.8 | 2.8 | 6 | 2.3 | 2.4 | 5 |
| 4 | 2.7 | 2.7 | 6 | 2.2 | 2.2 | 4 |
| 10 | 2.7 | 2.7 | 6 | 1.9 | 2.0 | 3 |
| 21 | 2.8 | 2.8 | 7 | 2.4 | 2.4 | 5 |
| 46 | 2.9 | 2.9 | 7 | 3.1 | 3.1 | 8 |
| 100 | 2.8 | 2.7 | 6 | 2.5 | 2.6 | 5 |
| 215 | 2.8 | 2.8 | 7 | 3.2 | 3.2 | 9 |
| 464 | 2.5 | 2.5 | 5 | 2.7 | 2.7 | 6 |
| 1000 | 2.8 | 2.8 | 6 | 2.6 | 2.6 | 5 |

## Discussion

We showed that, while the little auk social call is not a particularly loud signal (maximum 63 dBpeak re 20 μPa for both call types; compared to the loudest species reaching 140–150 dBpeak re 20 μPa in air; see e.g. Jakobsen *et al*. 2021), it is capable of carrying individual information over large distances. Calls could be classified to callers with very similar reliability independently of the distance, and well over the likely active space of the signal.

The *classic* call is the longest and most complex of the little auk repertoire (Osiecka *et al*. 2023a). Conspicuous signals are generally thought to have evolved for two main reasons: signalling quality and signal efficacy (Dawkins and Guilford 1997). The *classic* call certainly matches the latter description, maintaining its carrying capacity over distance. Similarly to other seabird vocalisations (e.g. Jones *et al*. 1987, Aubin *et al*. 2000, Curé *et al*. 2016, Baciadonna *et al*. 2021), little auk calls are reliable ‘self-reporting signals’ (Smith and Harper 1995), i.e. they provide information about the signaller. They carry cues to identity, notably in their fundamental frequency (Osiecka *et al*. 2024a), which has been shown here to travel over 1 km. However, little auks’ vocal identity can be somehow diluted when considering some parameters, since nesting partners match certain parameters of their calls, such as formant dispersion (Osiecka *et al*. 2023b). From a propagation perspective, as higher frequency formants are attenuated earlier on (see Fig. 6), suggesting that partners’ vocalisations become less similar with the distance, this may result in a seemingly increasing individual information content as the *classic* call travels further and further.

Long, complex signals can be used in long-distance communication in both humans (Seifart *et al*. 2018), and non-human animals (e.g. Dawkins and Guilford 1997, Luo *et al*. 2013, Larsen 2020). One aspect of the *classic* call that we did not investigate here is individuality coding within the temporal patterning of the call’s syllables – which in fact holds some of the parameters with the highest potential for individuality coding (Osiecka *et al*. 2024a). This was omitted due to the very heavy workload required to extract this information from such a large dataset. Nevertheless, the fact that strong individuality was retained even when excluding those parameters supports the notion that this call type is “designed” to facilitate efficient communication of identity. Adding the temporal information would very likely further increase the information content measured here, and improve clustering efficiency.

On the other hand, brevity often characterises short-distance communication (Luo *et al*. 2013). The *single* call is a very short, simple signal. While the classification efficiency of this call was essentially similar over distance, its information content dropped by roughly a half within the first two meters of propagation. This may suggest that even though this call type could be correctly classified over big distances, its primary role may lie more within short-range communication, i.e. to encode private information (Larsen 2020).

Of course, retaining information over long distances does not automatically translate into eliciting behavioural reactions to it. For instance, the corncrake *Crex crex,* whose calls carry cues to individuality over long distances (Ręk and Osiejuk 2011), but only result in response at behaviourally relevant distances (Ręk and Osiejuk 2010). However, the little auk’s Umwelt is very different of this of a corncrake, and such efficient long-distance communication could prove particularly useful. For instance, vocalisations could facilitate important aspects of a little auk’s life that might require individual recognition at long-distances, such as communication at foraging grounds, locating one’s neighbours or partner after migration, or even facilitating migratory behaviours. Dedicated studies are necessary to understand whether and how sound might play a role in these behaviours.

Long distance communication in the atmosphere is more likely to occur in environments with less physical constraints for sound transmission. For example, open habitats, such as the Arctic tundra or the sea degrade acoustic signals less than closed habitats (Hardt and Benedict 2020). However, acoustic communication in the atmosphere is also constrained by a number of factors contributing to signal attenuation, such as air humidity, temperature, and pressure (Wiley and Richards 1978). In response to this, animal signals can evolve to match the optimal frequency ranges for sound communication within their environments. The acoustic adaptation hypothesis (i.e. the notion that the vocal signal of a species will follow their habitat structure, e.g. open/closed) finds only some evidence and only in certain groups (Ey and Fischer 2009, Freitas *et al*. preprint), and a better match between signal properties and the environment can possibly be found at more local scales (as is e.g. the hooded crow, *Corvus cornix;* Jensen *et al*. 2008). While the fact that the Arctic tundra, as an open, humid habitat provides excellent conditions for sound propagation is not surprising, the reliability of information transmission over such distances is noteworthy.

So how far away from each other can two little auks be and still recognise the other’s voice, or react to it? This remains unknown, as here, we could only show that the signals themselves can be reliably classified to a sender at least up to the putative physiological hearing threshold, i.e. over the likely effective distance. This should be considered in the frame of information *content* and *transfer*, and not *meaning* (Weaver’s Levels A and B of communication problems; Shannon and Weaver 1949). That is, we cannot and do not intend to suggest to what level do little auks actually decode those transmitted signals and attribute them to individuals they know and recognise. Playback experiments in controlled conditions would be the only way to understand whether and how far away do little auks actually respond to such signals.

### Caveats and issues

This study, of course, comes with a number of limitations. While we are confident that the propagation model employing spherical spreading is appropriate for the studied vocalisation (uttered at great heights in an open habitat), it is necessarily simplified and does not correct for subtle changes to air layer densities, wind speed, or topography (see Guibard *et al*. 2022 for a brilliant model of ground surface communication in mountain habitats).

It is also likely that this study underestimates the sound pressure levels of the calls: because we had to be sure of the identity of the caller, we could only use calls produces within the nest, where we can control for it. However, calls uttered in open spaces are likely to have a higher amplitude than those produced in the nest, simply because of the increased noise outside (i.e., the Lombard effect, see e.g. Brumm & Zollinger 2013). Therefore, our study likely underestimated the real-life sound pressure levels, and therefore the active space, of these vocalisations when they are produced in flight. While this is unfortunate, we feel more confident reporting under-than over-estimated values.

Perhaps the biggest issue encountered here is that the recording distance (10 cm) falls within the near field of the lower frequency components of the calls – that is, the distance at which the soundwave is not yet fully developed, and might therefore behave differently (Larsen & Wahlberg 2019). Again, this is because recording the birds inside the nest was the only feasible way of obtaining repeated recordings of known individuals in the field. While the near field should not be an issue for the higher frequency components of the little auk calls, we acknowledge that the recorded properties of the lower components might not fully reflect the actual sound properties at larger distances.

This study also provides a somewhat idealised version of signal transmission, free of environmental noise and wind that surely interfere with the signal in real life: from other birds calling to glaciers calving, there are plenty of other sounds masking the little auk signals. Sadly, we were unable to perform propagation experiments due to the great heights and distances involved, and we acknowledge the importance of the local excess attenuation that was hence unaccounted for (see e.g. Jensen *et al*. 2008 and Guibard *et al*. 2022 for theoretical propagation models confirmed experimentally). Nevertheless, taking into account that the purpose of this study was to investigate the theoretical distance those signals can travel – and not how the animals use or perceive them – we believe that this framework still provides useful insights into the acoustic world of this little understood seabird.

## Conclusions

We found that the carrying capacity of the little auk social call does not decrease with distances over the likely behaviourally useful range. While these results do not indicate whether this information is actually perceived by the animals, this study suggests that vocal communication is likely used in long-distance communication, and can potentially facilitate important social interactions.

## Acknowledgements

Many thanks to the Institute of Geophysics of the Polish Academy of Sciences for providing access to long-term meteorological data, all the persons involved in fieldwork during data collection, Dariusz Jakubas for providing GPS readings of the Hornsund colonies, Damaris Riedner for advise on calibration methods, and to Ole Næsbye Larsen for priceless feedback on the early versions of this work.

This study was funded by grants awarded to the following authors: KWJ: grant no. 2017/25/B/NZ8/01417 funded by The National Science Centre (NCN), AO: University of Gdańsk Grants no. MN 539-D050-B853-21 and UGFirst 533-0C20-GF12-22.

## Authors’ contributions

AO: conceptualisation, funding acquisition, formal analysis – sound analysis, data analysis, visualisation, writing – original draft, review and editing. PB: conceptualisation, formal analysis – data analysis, visualisation, writing – original draft, review and editing. EFB: supervision, writing – review and editing. KWJ: funding acquisition, fieldwork, supervision, writing – review and editing.

## Ethics statement

This study used previously published data in theoretical models, and did not involve direct contact with the animals. Fieldwork involved in previous data collection was performed under permit from the Governor of Svalbard (20/00373-2), following the Association for the Study of Animal Behaviour’s guidelines for animal research.

## Data availability statement

Raw data and full codes generated in this study are available at https://osf.io/esbdj/?view_only=98d5eca13d894a94b9df1c1a091661fd. Audio files obtained from Osiecka *et al*. 2024a (https://doi.org/10.1016/j.anbehav.2024.02.009), https://osf.io/q9xhd/?view_only=2b8dd1470996468ea8f961d35070d1e5

## Supplementary Materials

**Supplementary Table 1.** **Kaiser-Meyer-Oklin factor adequacy: the overall KMO value for the dataset is middling for both call types, and data suitable for factor analysis.**

| Raw variable | *classic call* | *single call* |
| --- | --- | --- |
| Duration | 0.59 | 0.54 |
| AM Env Dep mean | 0.74 | 0.51 |
| AM Env Freq mean | 0.66 | 0.50 |
| AM Ms Freq mean | 0.71 | 0.76 |
| Ampl mean | 0.91 | 0.71 |
| CPP mean | 0.83 | 0.64 |
| Dom mean | 0.53 | 0.63 |
| FM Dep mean | 0.73 | 0.75 |
| Peak Freq mean | 0.83 | 0.91 |
| Pitch mean | 0.55 | 0.66 |
| Q25% | 0.81 | 0.79 |
| Q50% | 0.78 | 0.84 |
| Q75% | 0.76 | 0.72 |
| Spec Centroid mean | 0.73 | 0.71 |
| Spec Slope mean | 0.66 | 0.90 |
| **Overall KMO** | **0.75** | **0.74** |

**Supplementary Table 2.** **Principal Components Analysis: eigenvalues and proportion of variance.**

|  | *classic call* |  |  | *single call* |  |  |
| --- | --- | --- | --- | --- | --- | --- |
|  | **Eigenvalue** | **Proportion**<br>**of variance** | **Cumulative**<br>**proportion** | **Eigenvalue** | **Proportion**<br>**of variance** | **Cumulative**<br>**proportion** |
| **PC1** | 2.20 | 0.32 | 0.32 | 2.31 | 0.36 | 0.36 |
| **PC2** | 1.55 | 0.16 | 0.48 | 1.67 | 0.19 | 0.54 |
| **PC3** | 1.30 | 0.11 | 0.60 | 1.17 | 0.09 | 0.63 |
| **PC4** | 1.09 | 0.08 | 0.68 | 1.09 | 0.08 | 0.71 |
| **PC5** | 1.01 | 0.07 | 0.74 | 1.01 | 0.07 | 0.78 |
| **PC6** | 0.98 | 0.06 | 0.81 | 0.95 | 0.06 | 0.84 |
| **PC7** | 0.88 | 0.05 | 0.86 | 0.87 | 0.05 | 0.89 |
| **PC8** | 0.74 | 0.04 | 0.90 | 0.76 | 0.04 | 0.93 |
| **PC9** | 0.70 | 0.03 | 0.93 | 0.58 | 0.02 | 0.95 |
| **PC10** | 0.60 | 0.02 | 0.95 | 0.48 | 0.02 | 0.97 |
| **PC11** | 0.45 | 0.02 | 0.97 | 0.40 | 0.01 | 0.98 |
| **PC12** | 0.44 | 0.01 | 0.98 | 0.37 | 0.01 | 0.99 |
| **PC13** | 0.43 | 0.01 | 0.99 | 0.33 | 0.01 | 1.00 |
| **PC14** | 0.31 | 0.01 | 1.00 | 0.27 | 0.01 | 1.00 |
| **PC15** | 0.12 | 0.00 | 1.00 | 0.10 | 0.01 | 1.00 |

**Supplementary Table 3.** **Principal Components Analysis: contributions of raw acoustic parameters to the first five principal components of both call types**

|  | *classic call* |  |  |  | *single call* |  |  |  |
| --- | --- | --- | --- | --- | --- | --- | --- | --- |
| **Raw variable** | **PC1** | **PC2** | **PC3** | **PC4** | **PC1** | **PC2** | **PC3** | **PC4** |
| Duration | -0.10 | -0.50 | -0.30 | -0.01 | 0.28 | -0.72 | -0.34 | 0.23 |
| AM Env Dep mean | 0.01 | 0.66 | 0.42 | 0.15 | -0.26 | 0.65 | 0.62 | -0.11 |
| AM Env Freq mean | -0.31 | 0.19 | 0.42 | -0.64 | 0.35 | -0.08 | -0.56 | -0.52 |
| AM Ms Freq mean | -0.05 | -0.47 | -0.37 | 0.21 | -0.17 | 0.17 | -0.56 | 0.06 |
| Ampl mean | -0.04 | -0.06 | 0.00 | 0.27 | 0.11 | 0.47 | -0.33 | 0.48 |
| CPP mean | -0.68 | 0.20 | -0.03 | 0.54 | 0.42 | 0.53 | -0.30 | 0.37 |
| Dom mean | -0.28 | 0.44 | -0.71 | -0.23 | 0.24 | -0.70 | 0.20 | 0.43 |
| FM Dep mean | 0.04 | -0.73 | 0.05 | -0.36 | 0.30 | -0.30 | -0.21 | -0.25 |
| Peak Freq mean | -0.87 | -0.20 | 0.14 | -0.15 | 0.85 | -0.06 | 0.24 | -0.19 |
| Pitch mean | -0.31 | 0.42 | -0.74 | -0.26 | 0.41 | -0.61 | 0.39 | 0.25 |
| Q25% | -0.83 | 0.27 | -0.01 | 0.12 | 0.81 | -0.11 | 0.08 | -0.29 |
| Q50% | -0.78 | 0.26 | 0.11 | -0.19 | 0.93 | 0.02 | 0.17 | -0.15 |
| Q75% | -0.91 | -0.22 | 0.12 | -0.00 | 0.88 | 0.24 | 0.05 | 0.11 |
| Spec Centroid mean | -0.95 | -0.15 | 0.08 | 0.09 | 0.95 | 0.22 | 0.04 | 0.09 |
| Spec Slope mean | -0.58 | -0.51 | -0.03 | 0.07 | 0.79 | 0.47 | -0.10 | 0.15 |

